# Vaccination of pregnant mares with mRNA encoding a GPI-anchored form of the virulence-associated protein A of *Rhodococcus equi* induces strong IgG responses that are transferred to their foals

**DOI:** 10.64898/2026.10.26.762090

**Authors:** Noah D. Cohen, Nathan M. Canaday, Athos S. de Oliveira, Shelby Z. Wehmeyer, Cassidi N. Velez, Macy J. Fox, Maria T. Sanabria, M.H. Jesie Frisbie, Kerstin K. Landrock, Karin E.R. Borba, Ashley L. Jenkins, Angela I. Bordin, Jeroen B.K. Pollet

## Abstract

The bacterium *Rhodococcus equi* is a common cause of severe pneumonia in foals worldwide. Foals are exposed to *R. equi* from birth and are most susceptible in the first weeks of life. In North America, control measures are limited to transfusion to newborn foals of hyperimmune plasma whose potency is assessed by surpassing a threshold of IgG against the virulence-associated protein A (VapA) of *R. equi*. Here we report results of vaccination of pregnant mares with mRNA encoding a GPI-anchored virulence-associated protein A (VapA) of *R. equi* induced VapA-specific IgG subisotypes transferred to their foals during the 63-day period after birth. Pregnant mares were randomly assigned to receive either VapA mRNA vaccine (n=10) at 8 and 10 months of gestation or no vaccine (n=10). Serum was collected from mares at 8 months of gestation (before vaccination for immunized mares) and ≈24 hours after foaling and from foals at ages 1, 35, and 63 days. Mean relative concentrations of were low before immunization in all mares and did not change over time in unvaccinated mares; however, mean relative concentrations of anti-VapA IgG_1_, IgG_4/7_, and IgG_3/5_ of mares were significantly increased after VapA mRNA immunization and strongly correlated with the corresponding concentrations in their foals’ sera at age ≈24 hours. Concentrations of anti-VapA IgGs remained increased in foals of vaccinated mares compared with control foals for at least 35 days.

**IMPORTANCE:** Results of our study indicate that maternal vaccination of pregnant mares with a GPI-anchored VapA mRNA vaccine provides humoral immunity to foals when they are most susceptible to infection. These findings indicate that further evaluation of the efficacy of this vaccine to prevent pneumonia in foals infected with *R. equi* is warranted.

## INTRODUCTION

Pneumonia in foals caused by the bacterium *Rhodococcus equi* is a common cause of disease and death in foals worldwide.^1^ Strains of *R. equi* that are virulent in foals are widespread in the environment at horse breeding farms including soil, bedding, and air in pastures and stalls.^1–10^ Infection occurs by inhalation of aerosolized bacteria into the lungs of foals where virulent *R. equi* subsequently invade and replicate within alveolar macrophages.^1–3^ Rhodococcal pneumonia is a welfare concern for the equine industry because of the high cumulative incidence and severity of the disease at farms where the disease occurs recurrently.^1^ Although rigorous systematic economic impact studies are lacking, costs resulting from veterinary care, long-term treatment including labor, and mortality of some foals are high at affected farms.^1^ In addition to immediate costs, *R. equi* pneumonia can have a long-term detrimental effect on the equine industry by adversely reducing athletic performance.^11,12^

Virulent strains of *R. equi* carry a transferable virulence plasmid that is essential for intracellular replication within foal macrophages.^9,10^ Among the genes carried on the plasmid, *vapA* encodes virulence-associated protein A (VapA) which localizes to the bacterial surface and to the membrane of *R. equi*-containing vacuole, where it interferes with phagosome maturation and promotes a permissive compartment for bacterial replication in macrophages.^13–17^ VapA induces consistent humoral and cellular immune responses during natural infection and experimental challenge of foals. High anti-VapA antibody concentrations in foals are associated with lower odds of rhodococcal pneumonia.^1,18–28^

Because highly effective prevention of rhodococcal foal pneumonia is lacking,^1^ great need exists to develop an effective vaccine to protect foals against *R. equi*. We previously showed that an mRNA vaccine encoding VapA is immunogenic in neonatal foals when administered intramuscularly, using a construct in which the native bacterial signal peptide (SP) was replaced with an equine SP to achieve expression and secretion in equine cells.^29^ On the bacterium, however, VapA is retained at the cell surface by a mechanism that remains unsolved, although the N-terminus domain of the mature protein has been shown to contribute.^30^ For this study, we designed a new construct in which VapA is anchored to the outer side of the plasma membrane of the transfected cells through a C-terminal glycosylphosphatidylinositol (GPI) attachment signal, displaying the full mature VapA sequence in an extracellular orientation (hereafter referred to as gVapA mRNA).^31^ We hypothesized that GPI-mediated surface display of VapA would enhance VapA-specific antibody responses.

Vaccination of pregnant mares offers an opportunity to enhance colostral transfer of VapA-specific antibodies, providing passive protection during the early neonatal period when foals may encounter *R. equi* before developing vaccine-induced immunity^32,33^ and when foals are most susceptible to rhodococcal infection.^34^ This approach leverages the mare’s mature immune system and circumvents limitations associated with neonatal vaccination, including immune immaturity and interference by maternally derived antibodies. We therefore vaccinated pregnant mares with gVapA mRNA to characterize the induced VapA-specific IgG subisotype responses, assessed antibody transfer to their foals, and evaluated the persistence of those antibodies during the first 9 weeks after birth.

## MATERIALS AND METHODS

### Ethics statement

All methods were performed in accordance with relevant guidelines and regulations for animal use and laboratory practices (Texas A&M University Infectious Biohazard Committee IBC No. 2017-105; IACUC AUP No. 2023-0202 CA), including informed consent for serum samples collected from client-owned horses.

### Study animals

Twenty Quarter Horse mares and their foals at the *6666* Ranch in Guthrie, Texas were included in the study. Pregnant mares were randomly assigned to VapA mRNA-vaccinated (n=10) or unvaccinated (n=10) groups. Although our sample size for this initial evaluation of vaccine immunogenicity was largely determined by available funding, power calculations using the pwr package in R (Version 4.5.2, Foundation for Statistical Computing Vienna, Austria) indicated that with 10 mares in each group we had >95% power to detect a difference between 100% of vaccinated mares and 25% of control mares developing a >4-fold rise in optical density (OD) ratio of sample to positive control (**ODR**) at a significance level of 0.05. Randomization was performed using the randomizr package in R and a listing of pregnant mares available for the project. All foals included in the study were deemed to be healthy at age ≈24 hours based on findings of physical examination by farm veterinarians, results of CBC within reference ranges, and concentrations of immunoglobulin G (IgG) > 800 mg/dL. After blood was collected for IgG testing at age ≈24 hours, all foals were transfused with *R. equi* hyperimmune plasma from a single manufacturer (Mg Biologics). Foals were excluded from the study if they had health disorders (such as neonatal isoerythrolysis, diarrhea, or sepsis).

### Vaccine design and production

#### In vitro transcription of VapA mRNA

A mammalian codon-optimized DNA fragment encoding mature VapA of *R. equi* (GenBank: BAA04768) was synthesized by Twist Biosciences (San Francisco, CA). The fragment comprised, in order, a human albumin signal peptide, a FLAG tag, the VapA sequence, and the GPI attachment signal of horse placental alkaline phosphatase. This fragment was cloned into a plasmid backbone containing a T7 promoter compatible with CleanCap reagent, 5’ and 3’ untranslated sequences and a templated poly(A) tail of 100 to 110 nucleotides. Plasmids were linearized and used as template for in vitro transcription using CleanCap Reagent AG and m^1^ΨTP instead of rUTP (Trilink Biotechnologies, San Diego, CA), according to the manufacturer’s instructions. After DNAse I treatment, gVapA mRNA was purified with silica-based columns and had its integrity and purity evaluated by electrophoresis.

#### gVapA mRNA in vitro expression and antigen localization

For western immunoblotting (WB) and immunocytochemistry (IC), gVapA mRNA was transfected using Lipofectamine MessengerMax (Thermo Fisher Scientific) into mouse dendritic cell line DC.2.4 and primary equine bronchial fibroblasts (EBF) following the manufacturer’s instructions. Cells were harvested or stained 20 hours after transfection. For WB, VapA was detected using mouse monoclonal anti-FLAG M2 (MilliporeSigma, Cat# F3165, 1: 1000 dilution) as primary antibody and alkaline phosphatase-conjugated goat anti-mouse (KPL, Cat#5220-0357, 1:3000 dilution) as the secondary antibody. For intracellular staining, cells were fixed and permeabilized with Cytofix/Cytoperm buffer; for surface staining, cells were fixed with Cytofix buffer (BD, Franklin Lakes, NJ). Staining used Alexa Fluor 488-conjugated mouse anti-FLAG antibody (Thermo Fisher Scientific, Cat. MA1-142-A488, 1:250 dilution). Cell culture conditions and detailed WB and immunofluorescence protocols were performed as previously described.^29,35^

#### Formulating mRNA in lipid nanoparticles (LNPs)

The mRNA vaccine candidate was formulated into LNPs at a nitrogen-to-phosphate ratio of 4:1 using the NanoAssemblr Ignite instrument (Precision Nanosystems Inc., Vancouver, BC, Canada). The LNPs comprised the ionizable lipid ALC-0315, the helper phospholipid DSPC (1,2-distearoyl-sn-glycero-3-phosphocholine), cholesterol, and the PEGylated lipid ALC-0159 at an approximate molar ratio of 46.3:9.4:42.7:1.6, respectively. The RNA concentration and encapsulation efficiency of each lot were determined using the RiboGreen RNA Assay Kit (Invitrogen, Thermo Fisher Scientific, Waltham, MA, USA; catalog no. R11490). The mean LNP size and polydispersity index of each lot were measured by dynamic light scattering (DLS) (Malvern Panalytical, Malvern, United Kingdom). The formulations were diluted to an RNA concentration of 400 µg/mL with 1× PBS containing 8% sucrose and aliquoted into sterile serum vials at a fill volume of 1.2 mL. The vaccines were shipped to the farm on dry ice and stored at −20 °C until use.

### Immunization and sample collection

Mares in the vaccinated group were immunized intramuscularly (IM) at 8 and 10 months of gestation (**Figure 1**). Serum was collected for enzyme-linked immunosorbent assay (ELISA) testing from all mares at 8 months of gestation (before vaccination for the immunized mares), from all mares and their foals ≈24 hours after foaling, and from foals again at ages 35 and 63 days. Mares were monitored daily for evidence of adverse reactions to immunization by some of the authors (SZW, NMC, MJF, MHJF).

**Figure 1.**
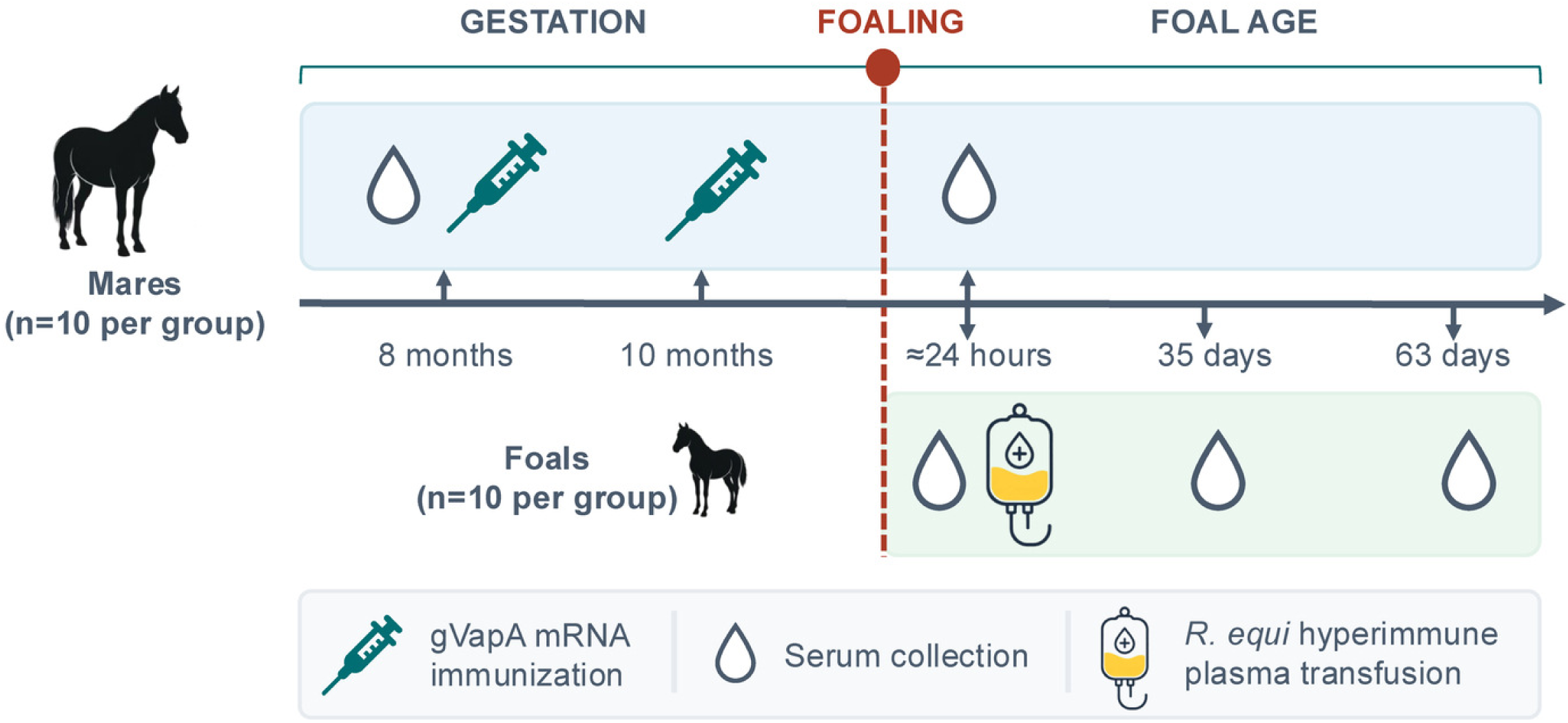
Schematic of study design. Mares in the vaccinated group (n=10) were immunized at 8 and 10 months with the GPI-anchored VapA mRNA vaccine, whereas control mares were not immunized. Serum was collected from mares at 8 months (prior to immunization for the vaccine group) and ≈24 hours after foaling. Serum was collected from foals at ages ≈24 hours (prior to transfusion with 2 L of *R. equi* hyperimmune plasma), 35 days, and 63 days.

### Anti-VapA IgG_1_, IgG_4/7_, and IgG_3/5_ ELISA

An indirect ELISA was used to determine the relative concentration of anti-VapA IgG_1_, IgG_4/7_, and IgG_3/5_ subisotypes. The following protocol was followed to ensure the accuracy of the assays. Before testing samples, we established an ELISA protocol that optimized the volumes and times for washing, incubation, and dilutions of test reagents, including blocking buffer and secondary antibodies. Quality control procedures were performed to verify the integrity of samples (i.e., no hemolysis) and reagents. Once a dilution protocol for samples and antigen concentration was established, we demonstrated that the intra-assay coefficient of variation (CV) was <15% and that the inter-assay CV was <15%. Serum samples were thawed prior to analysis. Immunoassay plates (Maxisorp ELISA plates, VWR Cat# 442404) were coated with recombinant VapA (custom-synthesized by the Protein Production, Characterization, and Molecular Interaction Laboratory, Texas A&M Institute of Biosciences and Technology) diluted with 1X coating buffer prepared from a 5X stock solution (Biolegend, Cat# 421701) to a concentration of 1 µg/mL VapA. Each well was coated with 100 µL of diluted VapA coating buffer and then incubated overnight at 4℃. Plates were washed 6 times with 1X PBS that contained 0.05% Tween-20. After washing, plates were blocked with 1% skim milk in 1X PBS (150 µL per well) for 1 hour at 37℃. The blocking buffer supernatant was discarded after incubation and serum samples (100 µL) were added in duplicate to the ELISA wells and incubated for 1 hour at 37℃. Serum samples used to test for anti-VapA antibodies for IgG_1_ were diluted 1/160, for IgG_4/7_ 1/10,240, and 1/320 for IgG_3/5_. Samples from each mare and their respective foals were tested on the same plate (i.e., mare at 8 months gestation, mare on day of foaling, foal at age 18 to 24 hours allowing for colostral antibody absorption). The positive control included in each ELISA plate was a serial dilution of serum from a donor horse that was hyperimmunized with *R. equi* generously provided by Mg Biologics, Inc. (Ames, IA), using 5 serial 2-fold dilutions starting at 1/80 for IgG_1_ and IgG_3/5_ and 1/2560 for IgG_4/7_. Dilution buffer was used for blank wells for both positive controls and serum samples. Following 6 washes, the subisotype specific detection antibody for IgG_1_ (1/4000 dilution; Bethyl Laboratories, catalog No. A70-124P), IgG_3/5_ (1/4000 dilution; Bethyl Laboratories, Cat# A70-105P), or IgG_4/7_ (1/8000 dilution; Bethyl Laboratories, Cat# A70-123P) was added to the respective plate, then incubated at room temperature for 1 hour. Following 6 washes, plates were treated with Sure Blue Reserve TMB Peroxidase Substrate (KPL; Cat# 5120-0081) and incubated for 20 minutes at approximately 21℃ in the dark. The reaction was stopped by adding a stop solution (0.18 M H_2_SO_4_), and optical densities (ODs) were measured at 450 nm using a microplate reader (Cytation 5; BioTek).

### Data analysis

All analyses were performed in R (Version 4.6.1) with significance set at P<.05. The outcome for analysis of anti-VapA IgG_1_, IgG_4/7_, or IgG_3/5_ in serum was the relative concentration represented by the ODR. The ODR was calculated as the mean OD of duplicate wells of the test sample (minus the mean of blanks for the plate) divided by the mean OD of the positive control at the same dilution as the test sample (minus the mean of the blanks). For inferential statistical analysis of the paired observations (i.e., before and after immunization) from mares or the serial samples in foals (i.e., ages 1, 35, and 63 days), data were analyzed using linear mixed-effects models in the nlme package with IgG subisotype ODRs as the outcome variable and vaccine group, sample-time, and their interaction modeled as fixed effects and individual mare or foal modeled as a random effect. Post hoc pairwise comparisons were made using the emmeans package implementing the Tukey method of adjustment. Correlation between VapA ODRs of mares and their foals was determined for each subisotype using Pearson’s correlation coefficients using the cor.test command in R. Correlation data were plotted with a linear regression model generated using the lm (linear model) command.

## RESULTS

### Expression and surface localization of gVapA

The gVapA mRNA encoded a 233-amino acid protein with a predicted molecular weight of 24.269 kDa (**Figure 2A**). Western immunoblotting of DC2.4 and EBF cells transfected with gVapA mRNA showed 2 bands close to the expected size that were absent from mock-transfected cells (**Figure 2B**). The lower band putatively corresponded to the signal peptide-cleaved form, and the upper band was consistent with the GPI-anchored form, in which the C-terminal signal was replaced by the glycolipid anchor and the apparent mass was increased.

**Figure 2.**
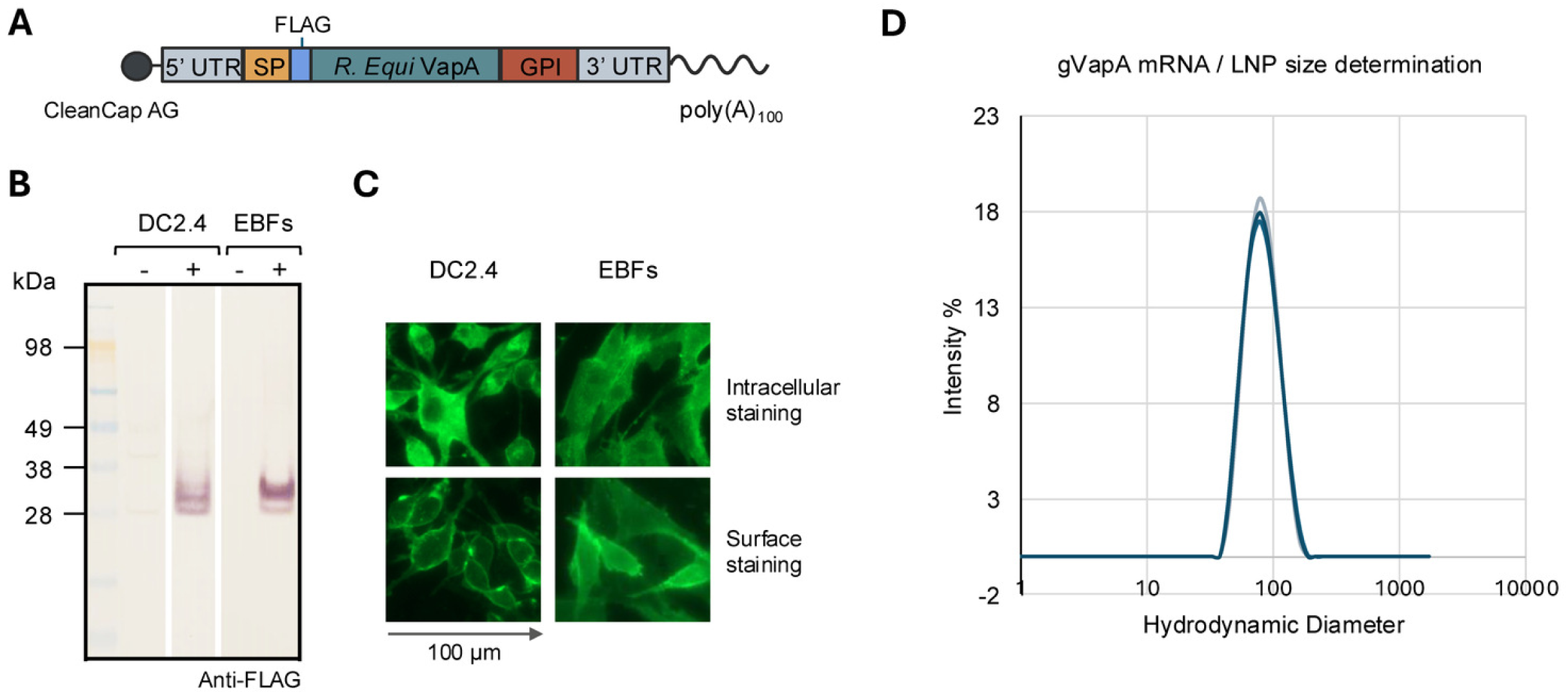
Design, in vitro expression, and formulation of the gVapA mRNA vaccine. **A.** Schematic of the gVapA mRNA construct. **B.** Western immunoblot of cell lysates (10 µg total protein per lane) from murine DC2.4 cells and primary equine bronchial fibroblasts (EBFs) transfected with gVapA mRNA (+) or mock-transfected (−), probed with anti-FLAG IgG as primary antibody. SeeBlue Plus2 pre-stained protein standard was used as size marker. **C.** Immunofluorescence of DC2.4 cells and EBFs transfected with gVapA mRNA. Cells were fixed and permeabilized (intracellular staining, upper row) or fixed without permeabilization (surface staining, lower row) and stained with Alexa Fluor 488-anti-FLAG. Cells were analyzed about 20 hours after transfection for WB and IC. **D.** Hydrodynamic diameter distribution of gVapA mRNA–lipid nanoparticles determined by dynamic light scattering; the trace shows 3 measurements of a representative preparation.

By IC, gVapA was detected in both cell lines under permeabilizing conditions (intracellular staining), with signal distributed throughout the cytoplasm in a reticular pattern, and under non-permeabilizing conditions (surface staining) where the signal outlined the plasma membrane (**Figure 2C**).

After formulation into LNPs, dynamic light scattering showed a single population of particles with a Z-average hydrodynamic mean diameter of 86 nm and a polydispersity index of 13% (**Figure 2D**). mRNA encapsulation was >85%.

### Serological responses of mares

Prior to immunization with the GPI-anchored VapA mRNA vaccine at approximately 8 months of gestation, baseline anti-virulence-associated protein A (VapA) ODRs for all IgG subisotypes (IgG_1_, IgG_4/7_, and IgG_3/5_) did not differ significantly between control mares (n=10) and mares allocated to the vaccinated group (n=10) (**Figures 3A, 3B, and 3C**; **Table 1**). By ≈24 hours post-foaling, anti-VapA ODRs for all 3 subisotypes were significantly (P<.0001) greater in vaccinated mares compared with either their pre-vaccination baseline values or control mares at either sampling time (**Figures 3A, 3B, and 3C**; **Table 1**). Importantly, control mares did not exhibit significant changes in anti-VapA IgG subisotype concentrations between sampling timepoints. No adverse effects were noted after vaccination at either timepoint in any mares. The estimated mean ODR for IgG_1_, IgG_4/7_, and IgG_3/5_ by time (i.e., 8 months of gestation and ≈24 hours post-foaling) and by vaccine group are summarized in Table 1.

**Figure 3.**
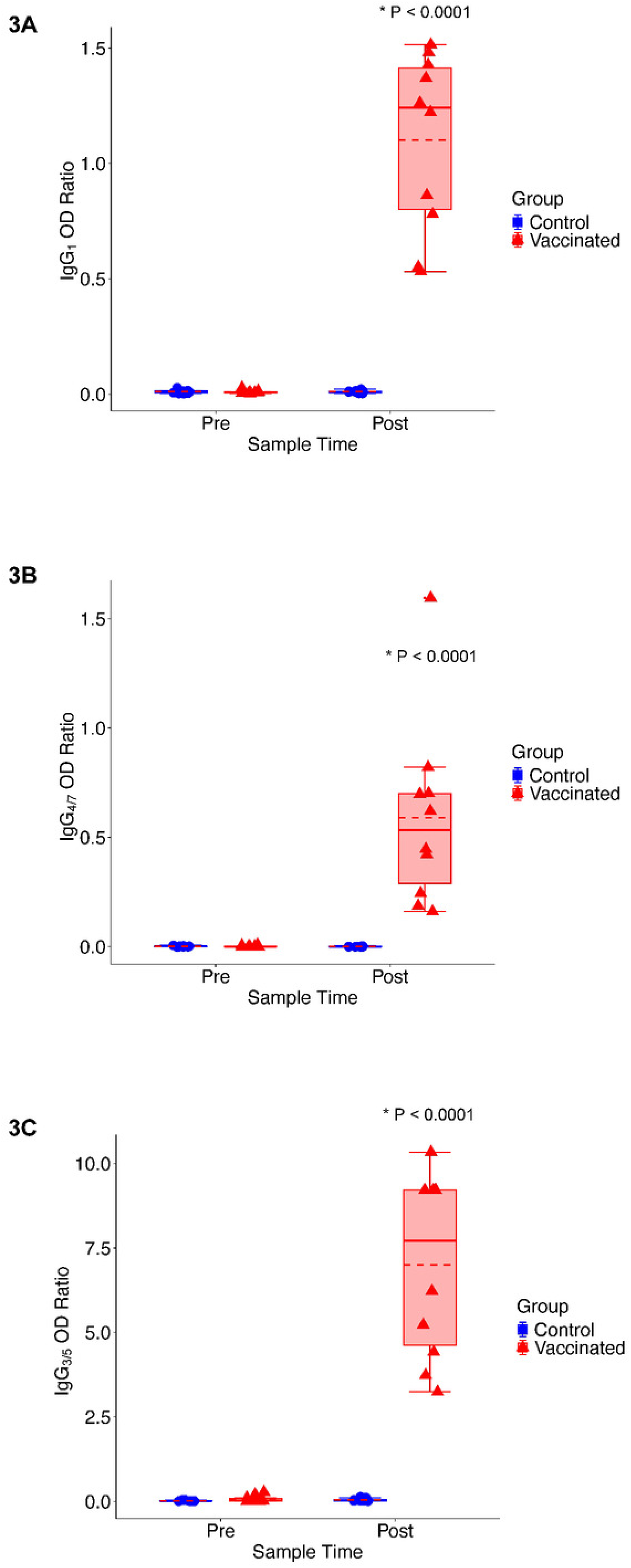
Relative concentrations of anti-VapA IgG_1_ (**A**), IgG_4/7_ (**B**), and IgG_3/5_ (**C**) in 10 mares vaccinated with VapA mRNA (red) and 10 control mares (blue) collected either at 8 months of gestation (Pre = prior to immunization in the vaccinated mares) or ≈24 hours after foaling (Post). Values were significantly (P<.0001) higher after foaling (Post) than before vaccination (Pre) in serum of mares vaccinated with VapA mRNA vaccine (n=10, red); concentrations from unvaccinated mares (n=10, blue) did not change between Pre and Post samples. Please see Table 1 for further details.

**Table 1.** Mean (95% CI) estimated optical density ratios (ODRs; i.e., relative concentration of sample to positive control) of anti-VapA Ig subisotypes in serum of mares by vaccine group and timepoint (Before = 8 months of gestation; After = 24 hours after foaling). Different superscripted numbers indicate significant (P<.0001) differences within rows.

|  |  | Control (n=10) |  | Vaccinate (n=10) |  |
| --- | --- | --- | --- | --- | --- |
|  | Subisotype | Before | After | Before | After |
|  | IgG<sub>1</sub> | 0.01 (-0.12 to 0.14)<sup>1</sup> | 0.01 (-0.12 to 0.14)<sup>1</sup> | 0.01 (-0.12 to 0.14)<sup>1</sup> | **1.10 (0.97 to 1.23)<sup>2</sup>** |
|  | IgG<sub>4/7</sub> | 0.00 (-0.14 to 0.14)<sup>1</sup> | 0.00 (-0.14 to 0.14)<sup>1</sup> | 0.00 (-0.14 to 0.14)<sup>1</sup> | **0.58 (0.45 to 0.73)<sup>2</sup>** |
|  | IgG<sub>3/5</sub> | 0.02 (-0.88 to 0.92)<sup>1</sup> | 0.05 (-0.86 to 0.95)<sup>1</sup> | 0.07 (-0.83 to 0.98)<sup>1</sup> | **7.00 (6.10 to 7.91)<sup>2</sup>** |

### Serological responses of foals

At ≈24 hours post-foaling and before transfusion of *R. equi* hyperimmune plasma, foals of vaccinated mares (n=10) had significantly higher serum concentrations of anti-VapA IgG subisotypes than foals of unvaccinated control mares (n=10) (**Figure 4A, 4B, and 4C**; **Table 2**). Strong and significant correlation was observed between mare and foal serum concentrations of anti-VapA IgG_1_ (Pearson’s r= 75; P=.0128), IgG_4/7_ (r=.85; P=.0017), and IgG_3/5_ (r=.74; P=.0153) when foals were ≈24 hours old (**Supplemental Materials**).

**Figure 4.**
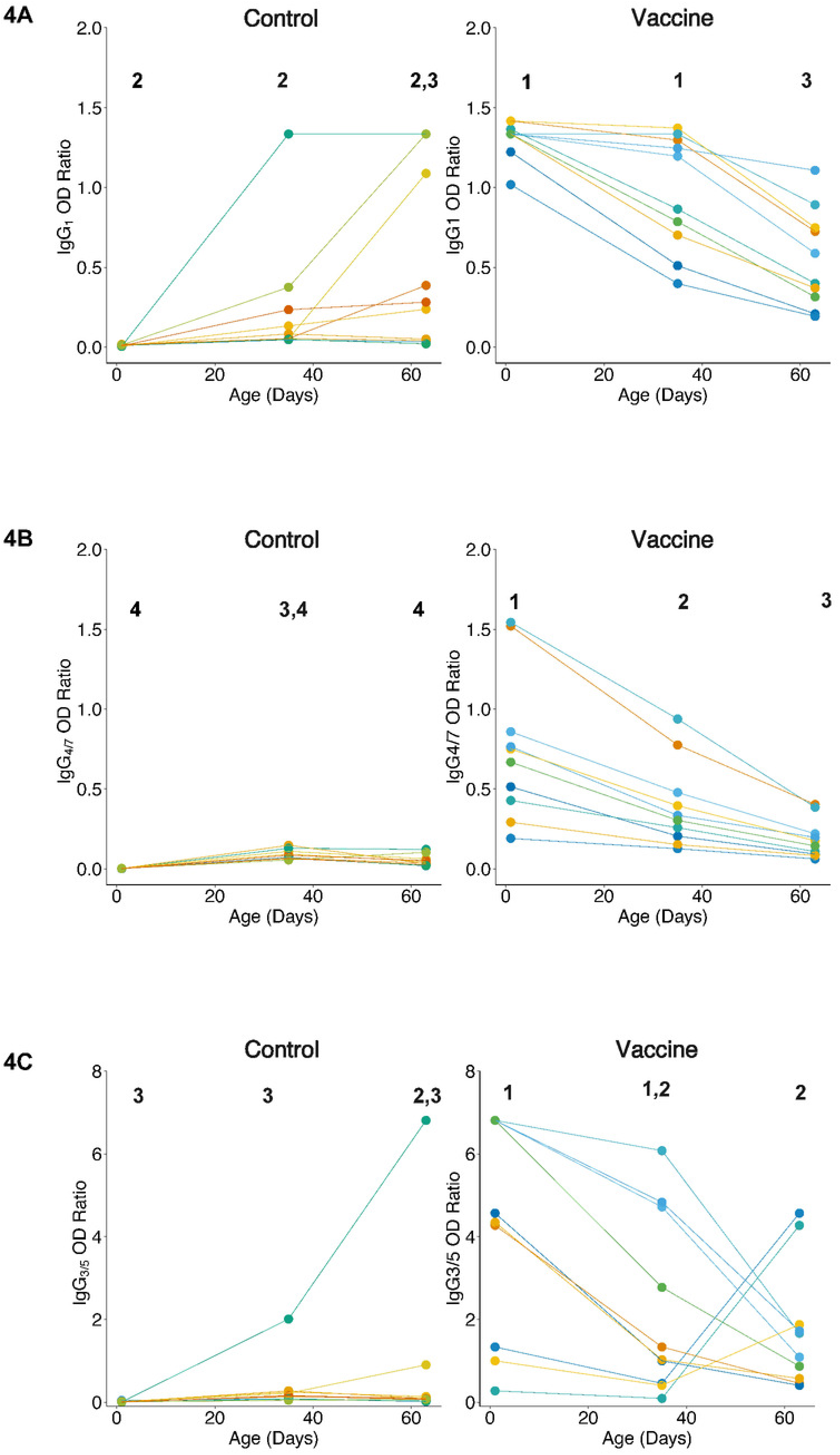
Relative concentrations of anti-VapA IgG_1_ (**A**), IgG_4/7_ (**B**), and IgG_3/5_ (**C**) by age in 10 foals of mares vaccinated with VapA mRNA (Vaccine; right) or 10 foals of unvaccinated control mares (left; Control). Serum samples were tested by indirect ELISA for anti-VapA antibodies at ages 1, 35, and 63 days. Bolded numbers denote significant (P<.05) pairwise differences in age and group for foals (summarized in Table 2).

**Table 2.** Mean (and 95% confidence intervals) of anti-VapA optical density ratios (ODRs) for IgG_1_ (a.), IgG_4/7_ (b.), and IgG_3/5_ (c.) by age for 10 foals of mares immunized with VapA mRNA or 10 foals of control mares not immunized with VapA mRNA. Values with different superscripted numbers differ significantly (P<0.05) within subisotype.

**a. IgG<sub>1</sub> ODRs**
| *Age (Days)* | *Vaccine Group* | *Control Group* |
| --- | --- | --- |
| **1** | 1.31 (1.08 to 1.54)<sup>1</sup> | 0.06 (-0.22 to 0.24)<sup>2</sup> |
| **35** | 0.97 (0.74 to 1.20)<sup>1</sup> | 0.24 (0.02 to 0.47)<sup>2</sup> |
| **63** | 0.55 (0.33 to 0.78)<sup>3</sup> | 0.48 (0.25 to 0.71)<sup>2,3</sup> |

**b. IgG<sub>4/7</sub> ODRs**
| *Age (Days)* | *Vaccine Group* | *Control Group* |
| --- | --- | --- |
| **1** | 0.75 (0.60 to 0.90)<sup>1</sup> | 0.00 (-0.15 to 0.15)<sup>4</sup> |
| **35** | 0.40 (0.25 to 0.55)<sup>2</sup> | 0.09 (-0.06 to 0.24)<sup>3,4</sup> |
| **63** | 0.19 (0.04 to 0.34)<sup>3</sup> | 0.05 (-0.10 to 0.20)<sup>3,4</sup> |

**c. IgG<sub>3/5</sub> ODRs**
| *Age (Days)* | *Vaccine Group* | *Control Group* |
| --- | --- | --- |
| **1** | 4.30 (3.14 to 5.47)<sup>1</sup> | 0.01 (-1.16 to 1.18)<sup>3</sup> |
| **35** | 2.27 (1.10 to 3.44)<sup>1,2</sup> | 0.34 (-0.83 to 1.51)<sup>3</sup> |
| **63** | 1.75 (0.58 to 2.92)<sup>2</sup> | 0.83 (-0.34 to 2.00)<sup>2,3</sup> |

All foals were transfused with 2 liters of *R. equi* hyperimmune plasma after serum was collected at age ≈24 hours. In foals of vaccinated mares, concentration of all 3 subisotypes remained significantly greater than control foals at age 35 days, but by 63 days the groups no longer differed significantly for any IgG subisotype (Table 2). Within the vaccinated group, the rate of decline differed among subisotypes: IgG_4/7_ decreased significantly between each consecutive sampling, whereas IgG_1_ and IgG_3/5_ did not differ significantly between ages 1 and 35 days and were significantly lower by 63 days. Three foals with the lowest serum concentrations of anti-VapA IgG_3/5_ at age 1 and 35 days increased in concentration between 35 and 63 days of age (**Figure 4C**). Relative concentrations of anti-VapA IgG subisotypes increased with age in some foals of unvaccinated mares, particularly for IgG_1_ and IgG_3/5_ (**Figure 4**).

## DISCUSSION

Intramuscular vaccination of broodmares during late gestation (months 8 and 10) with mRNA encoding GPI-anchored VapA safely produced strong maternal IgG responses against VapA: vaccinated mares had significantly higher post-foaling serum concentrations of anti-VapA IgG_1_, IgG_4/7_, and IgG_3/5_ compared with either their pre-vaccination levels or unvaccinated controls, and no adverse reactions to vaccines were noted. Foals born to vaccinated mares also had relatively high anti-VapA IgG_1_, IgG_4/7_, and IgG_3/5_ concentrations by 24 hours after birth that were strongly correlated with respective anti-VapA IgG subisotype concentrations in contemporaneous maternal serum samples, demonstrating effective passive transfer of vaccine-induced maternal antibodies. Although cell-mediated immunity and innate immunity are considered important in controlling *R. equi* infections in foals,^1^ increasing evidence indicates that antibodies can contribute to protection against intracellular pathogens.^36,37^

For both mares and foals, relative antibody concentrations were skewed toward IgG_1_ and IgG_3/5_ compared with IgG_4/7_: relative to serum concentrations of a hyperimmunized plasma donor horse, vaccination induced levels on average slightly higher than the positive control for IgG_1_ and several fold higher for IgG_3/5_, whereas the increase in IgG_4/7_ concentration was generally lower than that of the positive control (but still markedly higher than unvaccinated controls). It is possible this skewing reflects a difference in IgG subisotype distribution between immunization with mRNA and the proprietary immunization method used by the plasma manufacturer. The biological and clinical implications of this skewing are unknown. IgG_1_, IgG_3_, IgG_4_, and IgG_7_ have been demonstrated to have effector function such as fixing complement and stimulating respiratory burst by blood leukocytes.^38^ Higher relative concentration of anti-rhodococcal IgG_1_ has been associated with protection against *R. equi* infection in foals,^21,28,39–41^ and the fold-increase in serum anti-VapA antibody concentrations of adult horses after pulmonary infection with *R. equi* was greatest for IgG_1_.^42^ Thus, the IgG_1_ response observed following maternal vaccination might reflect skewing towards protective immunity.

The relative increase in concentration in vaccinated mares and their foals was greatest for IgG_3/5_. As noted above, IgG_3_ can mediate effector functions,^38^ but the role of this subisotype in *R. equi* infection is unclear. Because rhodococcal pneumonia is rarely if ever identified in immunocompetent adult horses despite widespread environmental exposure,^1–9^ adult horses are considered immune to *R. equi*. Because anti-VapA IgG_3/5_ has been observed to increase after pulmonary infection of adult horses,^42^ increased concentrations of IgG_3/5_ might be a correlate of protective immunity in foals. In foals, anti-VapA IgG_3/5_ in serum increased following natural infection with *R. equi* compared to foals from the same environment that did not develop pneumonia,^18^ and rose sooner after experimental infection with *R. equi*,^18^ suggesting that increased levels of serum IgG_3/5_ might reflect an effective immune response enabling foals to control infection with *R. equi*. Conversely, findings of higher anti-VapA IgG_3/5_ in foals affected with rhodococcal pneumonia relative to healthy foals from the same environment^18^ might reflect greater susceptibility to *R. equi* infection, such that the immune responses to our VapA mRNA vaccine might not be protective. Further study of the role of anti-VapA IgG_3/5_ in protection against *R. equi* is needed. Although the relative concentrations of IgG_4/7_ were lower than those of the other subisotypes, IgG_4/7_ is the subisotype in greatest serum concentration of horses and it too has effector functions that could help mediate protection against *R. equi*.^38^

The strong correlation of concentrations of anti-VapA IgG subisotypes between mares and their foals was consistent with maternal transfer of antibody to foals passively via suckling colostrum.^43^ Concentrations of anti-VapA IgG_1_, IgG_4/7_, and IgG_3/5_ generally decreased with age in foals of vaccinated mares, consistent with decay of maternally-transferred antibody.^43,44^ Nevertheless, anti-VapA IgG_3/5_ concentration increased between ages 35 and 63 days in 3 foals of vaccinated mares, presumably from natural exposure to *R. equi* in their environment. Notably, these 3 foals had the lowest concentrations of anti-VapA IgG_3/5_ at ages 1 and 35 days, indicating either that maternal antibody interference prevented increases in the other foals of vaccinated mares, or that these 3 foals were less likely to have been protected against *R. equi* infection than foals that had higher anti-VapA IgG_3/5_ concentrations. Several control foals had increased concentrations of anti-VapA IgG_1_ and 2 control foals had increased concentrations of anti-VapA IgG_3/5_. These increases were attributed to natural exposure to *R. equi* from their environment. The widespread natural exposure to *R. equi* where mares and foals are housed greatly complicates studies of immunity to rhodococcal foal pneumonia. Nevertheless, it appeared that maternal vaccination provided significantly higher concentrations of anti-VapA antibodies for at least the first 35 days after birth, a period when foals are most susceptible to infection.^34^

This study had limitations. The study population was relatively small and comprised a single breed at a single location. Immune responses were limited to assessing humoral responses; innate and T cell-mediated immune responses were not assessed. Immunity to *R. equi* is complex and incompletely understood, and it is possible if not probable that a combination of immune mechanisms including innate, cellular, and humoral immune responses contribute to protection.^1^ All foals were transfused with *R. equi* hyperimmune plasma according to standard practice for the farm. Transfusion did not impact results for foals at age 1 day because serum samples to test for anti-VapA antibodies were collected prior to transfusion. Given that the half-life of immunoglobulins in foals is approximately 3 weeks (21 days),^42,43^ it is unlikely that the observed differences in concentrations of IgG subisotypes in vaccinated and control foals at ages 35 and 63 days were attributable to transfusion. However, we cannot determine the impact of transfusion on immune responses to foals in this study. Notably, our results indicate that if the objective of transfusion is to increase anti-VapA antibody concentrations, maternal vaccination with mRNA might be more effective and more convenient than transfusing foals with 1 or 2 liters of *R. equi* hyperimmune plasma.^25^ The foremost limitation of our study is that we did not assess clinical efficacy of the vaccine in study foals. Efficacy studies will require a larger population of mares and foals and considerably more resources than were available for this project. Moreover, we considered it necessary to first determine the immunogenicity of our mRNA vaccine before considering the design and conduct of studies to assess vaccine efficacy. The strong immunogenicity of this VapA mRNA vaccine provides evidence to support future efficacy studies to address these limitations.

